# Limiting distribution of X-chromosomal coalescence times under first-cousin consanguineous mating

**DOI:** 10.1101/2022.05.05.489432

**Authors:** Daniel J. Cotter, Alissa L. Severson, Shai Carmi, Noah A. Rosenberg

## Abstract

By providing additional opportunities for coalescence within families, the presence of consanguineous unions in a population reduces coalescence times relative to non-consanguineous populations. First-cousin consanguinity can take one of six forms differing in the configuration of sexes in the pedigree of the male and female cousins who join in a consanguineous union: patrilateral parallel, patrilateral cross, matrilateral parallel, matrilateral cross, bilateral parallel, and bilateral cross. Considering populations with each of the six types of first-cousin consanguinity individually and a population with a mixture of the four unilateral types, we examine coalescent models of consanguinity. We previously computed, for first-cousin consanguinity models, the mean coalescence time for X-chromosomal loci and the limiting distribution of coalescence times for autosomal loci. Here, we use the separation-of-time-scales approach to obtain the limiting distribution of coalescence times for X-chromosomal loci. This limiting distribution has an instantaneous coalescence probability that depends on the probability that a union is consanguineous; lineages that do not coalesce instantaneously coalesce according to an exponential distribution. We study the effects on the coalescence time distribution of the type of first-cousin consanguinity, showing that patrilateral-parallel and patrilateral-cross consanguinity have no effect on X-chromosomal coalescence time distributions and that matrilateral-parallel consanguinity decreases coalescence times to a greater extent than does matrilateral-cross consanguinity.

## 1 Introduction

The phenomenon of consanguinity, in which unions occur between closely related individuals, is a form of population structure that can dramatically affect properties of genetic variation (Crow and Kimura, 1970; Jacquard, 1974). By increasing the probability that deleterious recessive variants appear in homozygous form, it contributes to the incidence of recessive disease (Bittles, 2001; Woods *et al*., 2006); recent studies suggest that it contributes to incidence of complex disease as well (Bittles and Black, 2010; Yengo *et al*., 2017; Ceballos *et al*., 2018; Johnson *et al*., 2018; Clark *et al*., 2019). Consanguinity is common in human populations, with some populations promoting consanguineous marriages as a cultural preference (Bittles, 2012; Romeo and Bittles, 2014; Sahoo *et al*., 2021).

The offspring of a consanguineous union are expected to possess large portions of their genomes shared between their two genomic copies, owing to the fact that an identical genomic segment can be inherited along both their maternal and paternal lines. For the loci contained in such segments, the two copies *coalesce* at a common ancestor relatively few generations in the past. At other locations, neither copy or only one copy traces to a recent shared ancestor, so that coalescence occurs only much farther back in the past. Indeed, empirical genetic studies have identified multiple populations in which individuals carry long runs of homozygosity (ROH), attributable in large part to consanguinity practices (McQuillan *et al*., 2008; Pemberton *et al*., 2012; Ceballos *et al*., 2018)

In typical coalescent-based models that investigate coalescence times for sets of lineages, diploid organisms are approximated by pairs of haploids independently drawn from a population (Hein *et al*., 2004; Wakeley, 2009). This modeling choice is unsuited to the study of consanguineous families, in which the two lineages in an individual can be highly dependent. Hence, explicitly diploid coalescent models have been devised for the study of coalescence in a setting of consanguinity. The earliest studies focused on selfing in plants (Pollak, 1987; Nordborg and Donnelly, 1997; Nordborg and Krone, 2002), an extreme form of “consanguinity” in which both parents of a diploid offspring are the same individual. Campbell (2015) extended diploid coalescent models to consider a monogamous mating model with sibling mating, computing mean coalescence times under the model. This approach was then extended by Severson *et al*. (2019) to consider mean coalescence times in a diploid model with *n*th-cousin mating, for arbitrary values of *n* and for superpositions of multiple levels of *n*th-cousin mating.

In an extension of the work of Severson *et al*. (2019), Severson *et al*. (2021) advanced beyond mean coalescence times to derive a full limiting distribution of coalescence times under superposition models of autosomal consanguinity, considering the limit as the population size grows large. A limitation of the work of Severson *et al*. (2019) and Severson *et al*. (2021), however, is that it does not distinguish between males and females in the mating model; all individuals are exchangeable. Hence, it cannot accommodate the variety of scenarios in which differences between males and females are salient. We have recently extended the method of Severson *et al*. (2019) to distinguish between males and females, evaluating mean coalescence times in a two-sex model, with a goal of evaluating the effect that consanguinity has on X-chromosomal coalescence times specifically (Cotter *et al*., 2021).

Here, we use the advance from Severson *et al*. (2021) to compute the full distribution of coalescence times under a diploid, two-sex consanguinity model (Cotter *et al*., 2021). Seeking to derive distributions of X-chromosomal coalescence times, we consider each of the six types of first-cousin consanguinity and a model that includes all four unilateral types in a single population. For each model, we evaluate the distribution of coalescence times for two lineages sampled from the same individual and for two lineages sampled from members of different mating pairs.

## 2 Methods

We adapt the models of Severson *et al*. (2019, 2021) and Cotter *et al*. (2021). We consider a constant-sized population of *N* diploid mating pairs. Individuals are sex-specific, the X chromosome is considered, and specified forms of consanguinity are allowed. Using a Markov chain, we track lineage pairs back in time until they coalesce.

To analyze the large-*N* limit of the model, we make use of the separation-of-time-scales approach introduced by Möhle (1998). This approach was used by Severson *et al*. (2021) to obtain the limiting distribution of coalescence times under their autosomal diploid model of consanguinity. In the approach from Möhle (1998), the limiting distribution of a Markov process with transition matrix Π_*N*_ is obtained by writing

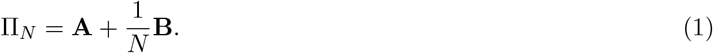

Here, **A** describes “fast” transitions that have nontrivial probability in a single generation, and **B** describes “slow” transitions that have very small probabilities in a single generation. As *N* → ∞, the fast transitions occur instantaneously, and the fast process can be described by an equilibrium distribution

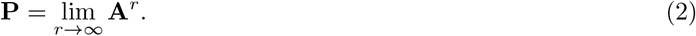

Rescaling *t* in units of *N* generations, as *N* → ∞, Π_*N*_ converges to a continuous-time process

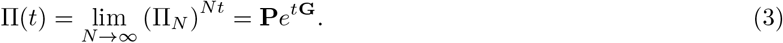

The rate matrix **G** satisfies **G** = **PBP**. Under Möhle’s theorem, the process converges to a continuous-time process with an instantaneous jump at time 0 that corresponds to the “fast” transitions.

As Severson *et al*. (2021) did with autosomal models, we apply the separation-of-time-scales approach to our models of consanguinity on the X chromosome (Cotter *et al*., 2021). We begin with the sib mating case and then consider each of the four types of unilateral first-cousin mating, the two cases of bilateral first-cousin mating, and a mixture of all four unilateral types in one model.

## 3 Results

### 3.1 Sibling mating

We consider *N* monogamous male–female mating pairs, a fraction *c*_0_ of which are sib mating pairs. Pairs of X-chromosomal lineages can be in one of six states (Figure 1): two lineages have already coalesced (state 0); two lineages are in a female (state 1); two lineages are in opposite individuals of a mating pair (state 2); two lineages are in two individuals in different mating pairs, where the two individuals are two males (state 3), a male and a female (state 4), or two females (state 5). Note that for the X chromosome, there is no state for two lineages in a male, as males contain only one X chromosome. We track the state of the process backward in time until it reaches the most recent common ancestor for a pair of lineages (that is, until state 0 is reached). We denote by *T*_*f*_, *U, V*_*mm*_, *V*_*mf*_, and *V*_*ff*_ the random coalescence time for pairs of lineages in states 1, 2, 3, 4, and 5, respectively.

**Figure 1:**
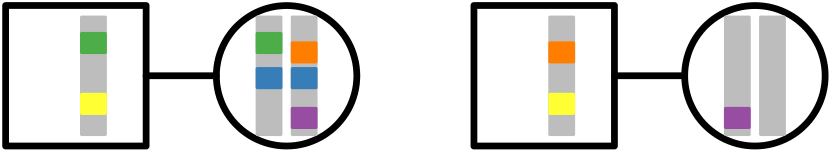
Five states for two lineages. Males are squares; females are circles. State 1: within a female (blue). State 2: in two individuals in a mating pair (green). State 3: in two males in different mating pairs (yellow). State 4: in a male and a female in different mating pairs (orange). State 5: in two females in different mating pairs (purple).

If two lineages are in state 0 (coalesced), they remain in state 0 with probability 1; this state is absorbing. If two lineages are in a female (state 1), in the previous generation they must have been in separate individuals in a mating pair (state 2) with probability 1. If two lineages are in separate individuals in a mating pair (state 2), the pair is a sib mating pair with probability *c*_0_. Given that the pair is a sib mating pair, the lineages transition to state 0 with probability 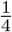, state 1 with probability 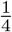, and state 2 with probability 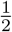. If the two lineages are not in a sib mating pair, an event with probability 1 − *c*_0_, then they transition to states 4 and 5 with equal probability 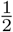.

For each of the states 3–5, because we pick parental mating pairs with replacement from the previous generation, the probability is 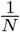 that the same mating pair is chosen. Thus, if two lineages are in state 3, and the pair are siblings (an event with probability 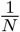), then the lineages transition to state 0 or state 1, each with probability 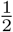.If the two lineages in state 3 do not have the same parental pair (probability 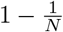), then they must transition to state 5 with probability 1. For state 4, if the two lineages are in siblings (probability 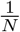), then they transition to state 0 with probability 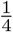, state 1 with probability 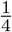, and state 2 with probability 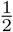. If the lineages are not from siblings (probability 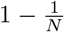), then they transition to state 4 or 5, each with probability 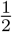. Finally, two lineages in state 5, conditional on being in siblings (probability 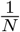), reach state 0 with probability 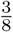, state 1 with probability 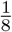, and state 2 with probability 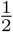. Conditional on not being in siblings (probability 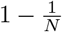), the lineages transition to state 3 with probability 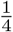, state 4 with probability 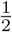, and state 5 with probability 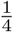.

Combining these transition probabilities, we can write the transition matrix as

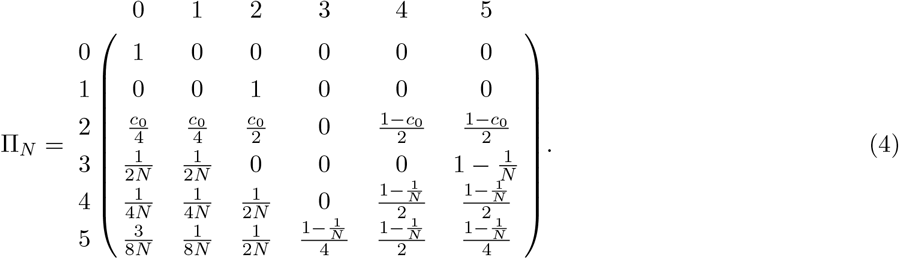

We can decompose Π_*N*_ (Eq. 4) into its fast and slow transitions, as in Eq. 1:

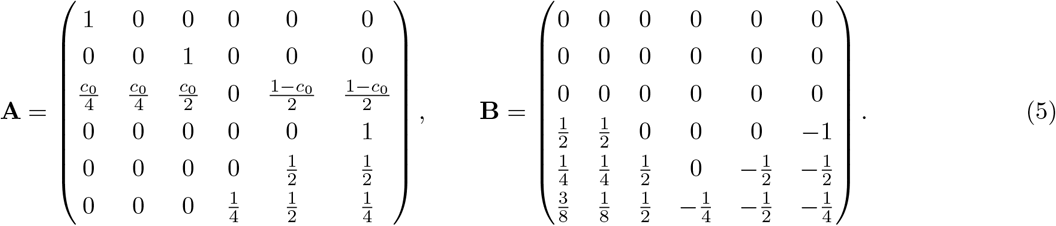

We first find the equilibrium distribution of the “fast” process, obtained by iterating transition matrix **A**. This calculation appears in Appendix A, producing

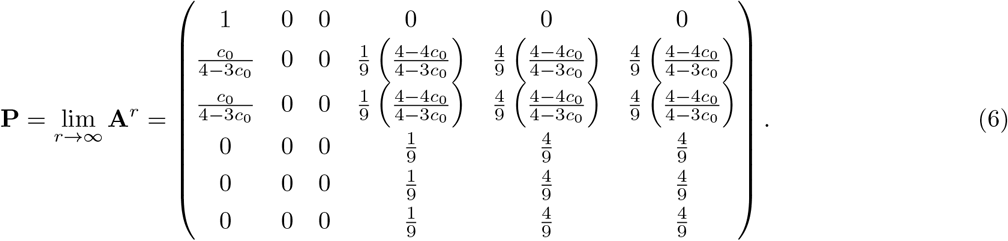

We then compute **G** = **PBP** and solve for the limiting process Π(*t*) using Eq. 3, obtaining the matrix exponential, *e*^*t***G**^, as in Appendix B. Converting *t* back into units of *N* generations, this gives

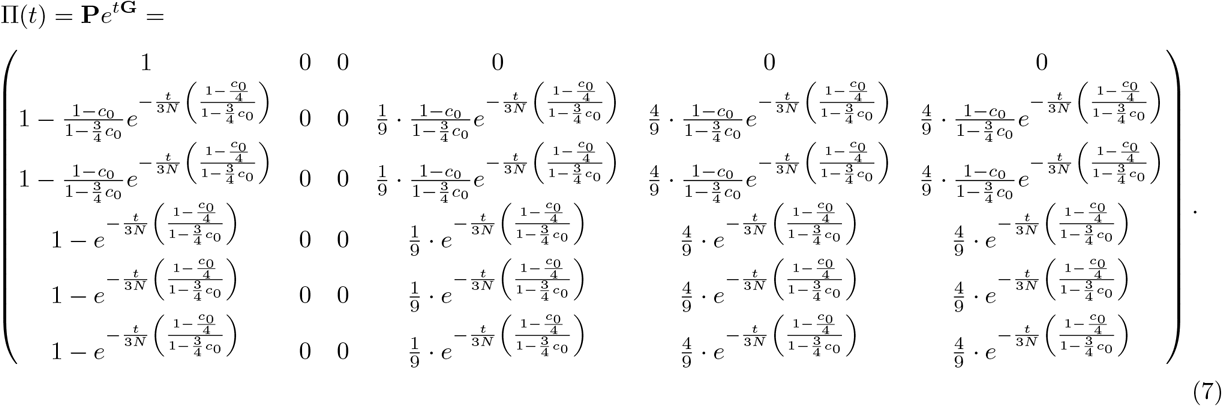

The first column of the matrix Π(*t*) represents the cumulative probability of coalescence in time less than or equal to *t* generations. States 1 and 2 have the same cumulative distribution, representing the coalescence time for two lineages *within* a female (note that state 2, two lineages in the two individuals in a mating pair, is always reached from state 1 after one step). States 3–5 have the same cumulative distribution, representing the coalescence time for two lineages in two distinct individuals. The cumulative distributions are

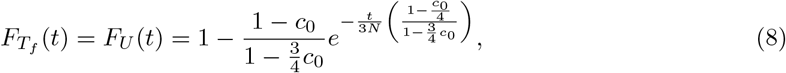

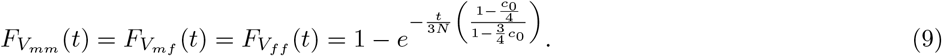

Computing the expectations of these distributions, recalling that for 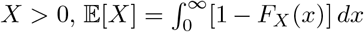, we find

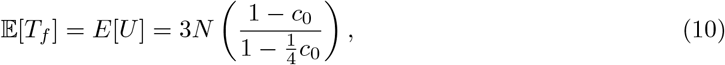

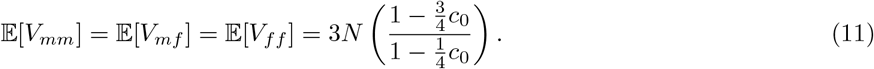

where Eqs. 10 and 11 are the same as Eqs. 25 and 26 from Cotter *et al*. (2021), obtained by first-step analysis.

Eqs. 8 and 9 are plotted in Figure 2. In the figure, we observe that the cumulative probability of coalescence increases with the consanguinity probability *c*_0_. For *c*_0_ = 0, 𝔼[*T*_*f*_] = 𝔼[*V*_*mf*_] = 3*N*, as there are three copies of the X chromosome in each mating pair in the population. For *c*_0_ *>* 0, 𝔼[*T*_*f*_] *<* 𝔼[*V*_*mf*_] due to the probability of consanguinity whenever the two lineages are already in the same mating pair.

**Figure 2:**
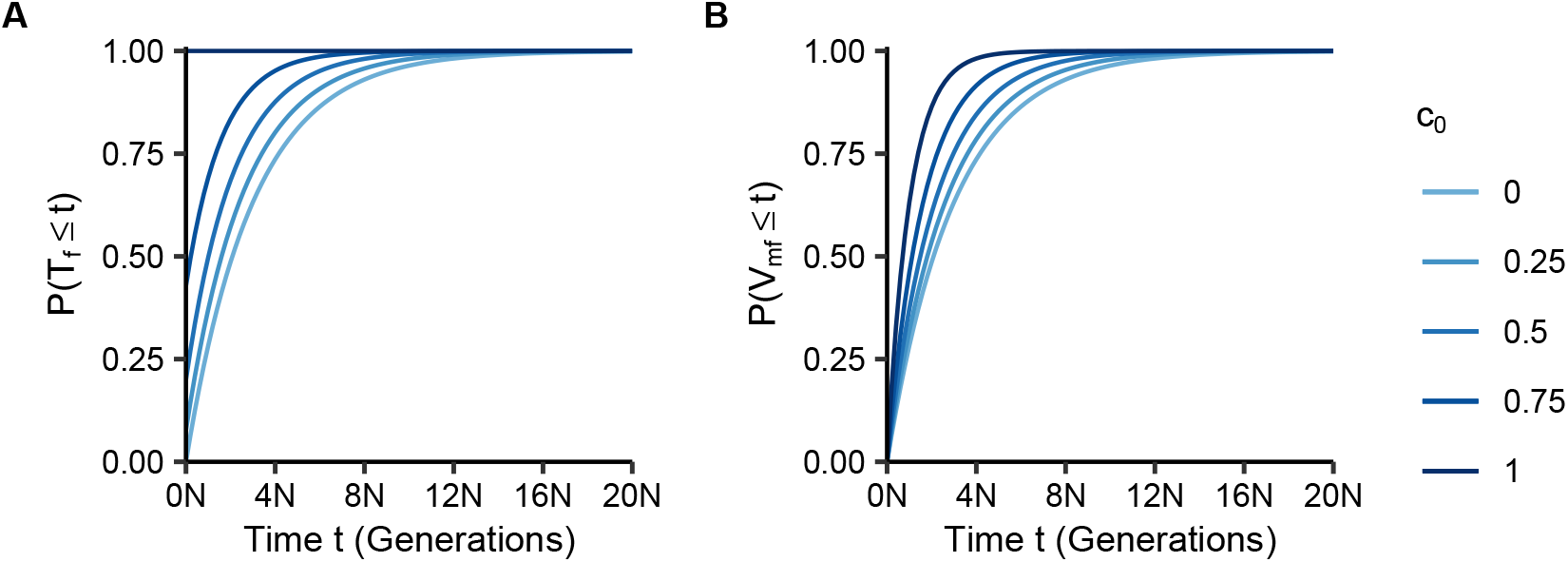
Cumulative distributions of coalescence times within (*T*_*f*_) and between (*V*_*mf*_) individuals as functions of the number of generations *t* and the fraction of sib mating pairs *c*_0_. **(A)** Within individuals, *P* (*T*_*f*_ ≤ *t*), Eq. 8. **(B)** Between individuals, *P* (*V*_*mf*_ ≤ *t*), Eq. 9.

### 3.2 First cousins

We next consider first-cousin consanguinity on the X chromosome. We separately calculate the limiting distributions of coalescence times for each of the four types of first-cousin consanguinity: patrilateral parallel, a union of a male with his father’s brother’s daughter; patrilateral cross, a union of a male with his father’s sister’s daughter; matrilateral parallel, a union of a male with mother’s sister’s daughter; and matrilateral cross, a union of a male with his mother’s brother’s daughter.

For each of these four types of first-cousin consanguinity, two lineages have seven possible states. State 0 is an absorbing state representing coalescence. State 1 is two lineages in a female. States 3–5 represent, as in the sibling case, two lineages that are in two individuals in *different* mating pairs, where the two individuals are two males (state 3), a male and a female (state 4), or two females (state 5).

Next, for pairs of lineages from the two individuals in a mating pair, we follow the model of a superposition of multiple mating levels from Severson *et al*. (2021), taking a special case of this approach. Under the superposition model, each state 2_*i*_, 0 ≤ *i* ≤ *n*, represents an ancestral state for two lineages from a mating pair. These ancestral states can be viewed as “holding states” that keep track of ancestral lineages of a mating pair in order to allow all possible *i*th-cousin levels of consanguinity up to *n*th cousins. As we restrict attention to first-cousin mating, we need only states 2_0_ and 2_1_ from Severson *et al*. (2021).

State 2_0_ represents two lineages in the two individuals in a mating pair. State 2_1_ represents two lineages in two individuals ancestral to the two individuals in a mating pair. Because, unlike Severson *et al*. (2021), we disallow sib mating, two lineages in state 2_0_ cannot coalesce (state 0), they cannot transition to the same individual (state 1), nor can they transition to two individuals in a mating pair (state 2_0_). Hence, lineages in 2_0_ must transition to 2_1_ (Figures 3 and 4).

**Figure 3:**
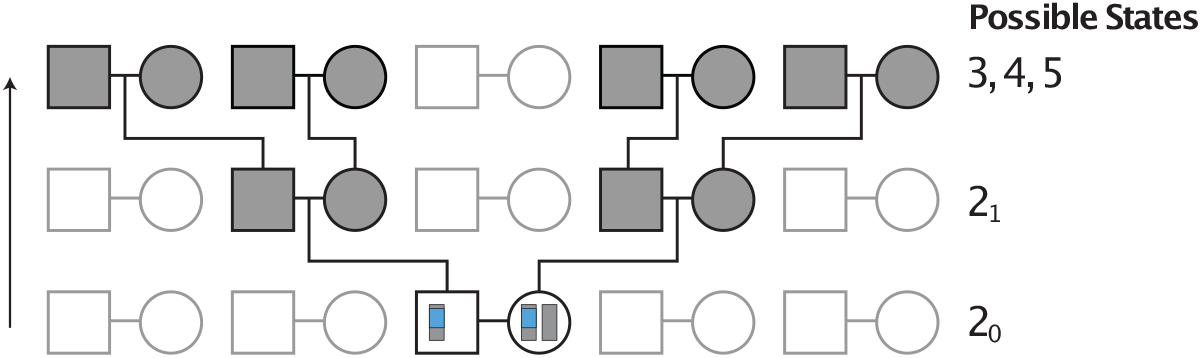
Example pedigree illustrating transitions from state 2_0_ in the absence of consanguinity. Considering a pair of lineages in a mating pair, depicted in blue, the process always immediately transitions to the holding state 2_1_ one generation in the past. From state 2_1_, the lineages transition to two separate mating pairs, and hence, to states 3, 4, or 5.

**Figure 4:**
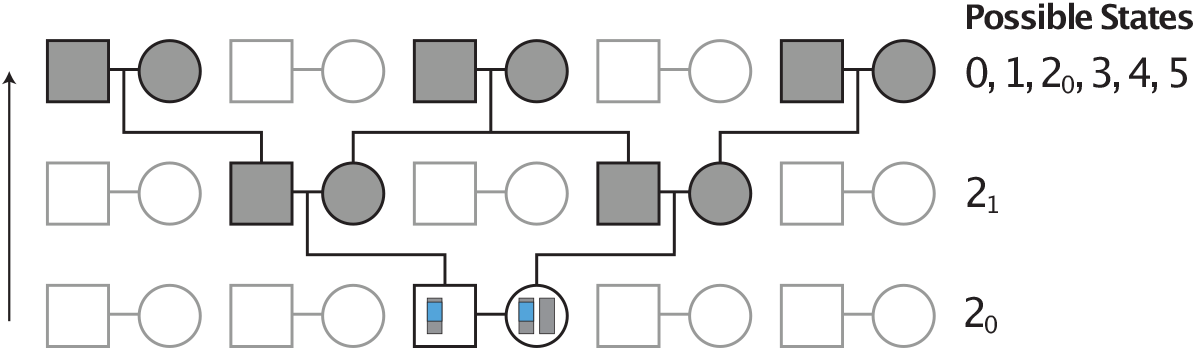
Example pedigree illustrating transitions from state 2_0_ in the presence of first-cousin consanguinity. Considering a pair of lineages in a mating pair, depicted in blue, the process always immediately transitions to the holding state 2_1_. From state 2_1_, the lineages can potentially transition to any of states 0, 1, 2_0_, 3, 4, 5, depending on the type of first-cousin consanguinity. Matrilateral-cross consanguinity is depicted.

In the absence of consanguinity, two lineages in state 2_1_ can transition only to states 3, 4, and 5 (Figure 3). With first-cousin consanguinity present (Figure 4), two lineages in state 2_1_ can also coalesce (state 0) or transition to two lineages in the same female (state 1) or to two lineages in opposite individuals in a mating pair (state 2_0_).

The transition matrix depends on the type of first-cousin consanguinity permitted. However, the type of consanguinity only affects transitions from state 2_1_. For all types of consanguinity, state 0 is an absorbing state. State 1, two lineages in the same female, always transitions to state 2_0_ because the two lineages must come from opposite individuals of the same mating pair. Because of the constraints we have placed on the process, state 2_0_ always transitions to state 2_1_. Finally, the transition probabilities from states 3, 4, and 5 follow the same pattern as given in the transition matrix in Eq. 4 (with state 2_0_ in place of state 2).

Below, we consider each of the four different types of first-cousin mating, two cases of bilateral first-cousin mating, and a mixture of the four unilateral types. In each case, we define the transitions that the process makes from state 2_1_, and we obtain the limiting distributions of coalescence times.

#### 3.2.1 Patrilateral parallel

In patrilateral parallel first-cousin consanguinity, a union occurs between a male and his father’s brother’s daughter. There is no way for the X-chromosomal lineages in the first-cousin mating pair to have originated from the shared grandparental pair, because X chromosomes are never transmitted from fathers to sons. Hence, irrespective of the fraction *c*_1_ in the population, lineages in state 2_1_ can only transition to states 3, 4, and 5.

In state 2_1_, one X chromosome in one of the parental pairs is always in a female (the parent of the male in state 2_0_). The probability is then 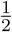 that this X chromosome is in a male one generation ancestral to 2_1_ and 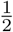 that it is in a female. The other X chromosome in state 2_1_, located in a parent of the female in state 2_0_, can be in a male or female, with equal probability. Hence, one generation ancestral to 2_1_, this X chromosome is in a female with probability 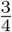 and in a male with probability 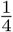. We can multiply probabilities for the two separate X chromosomes to obtain transition probabilities from state 2_1_. In particular, the two lineages will be in two separate males one generation previously (state 3) with probability 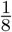. They will be in a male and a female (state 4) with probability 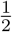. They will be in two separate females (state 5) with probability 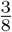.

The transition matrix is:

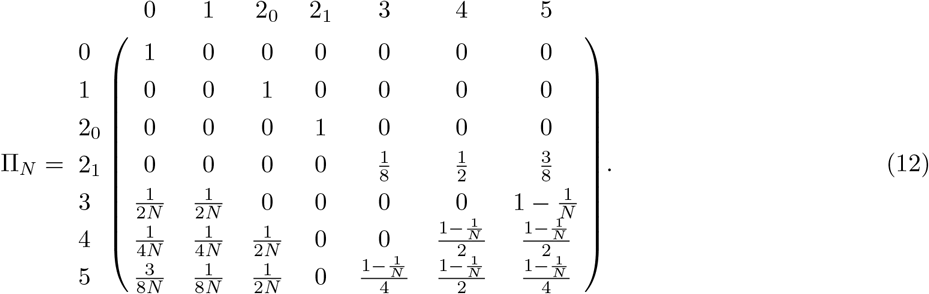

As with the sibling case, we can decompose the transitions into “fast” and “slow” transitions (Eq. 1):

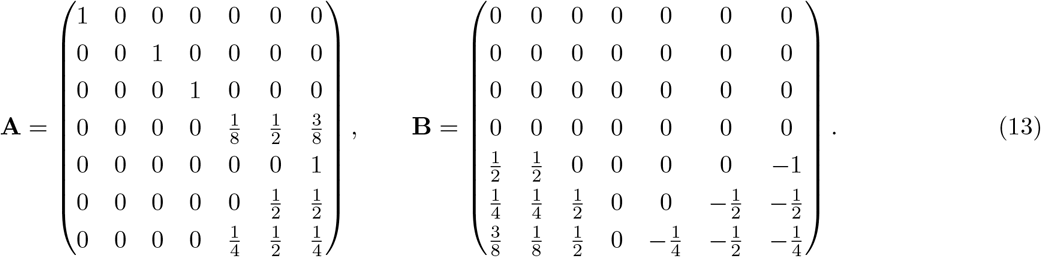

We next solve for the limiting distribution of the fast transition matrix **A** using the method of Appendix A,

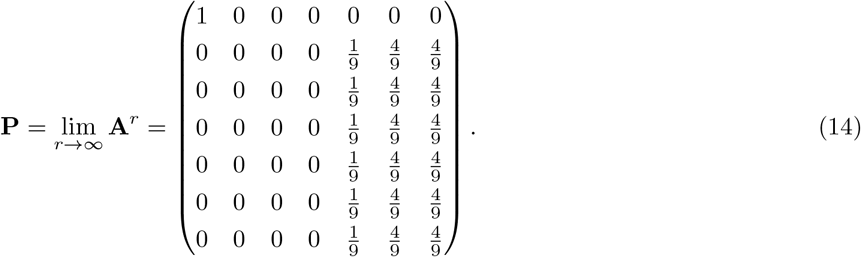

Recalling **G** = **PBP**, we solve for the limit Π(*t*) as in the sibling mating case, using Eq. 3, calculating the matrix exponential, *e*^*t***G**^, as in Appendix B. We then convert *t* back into units of generations *N*. This step gives

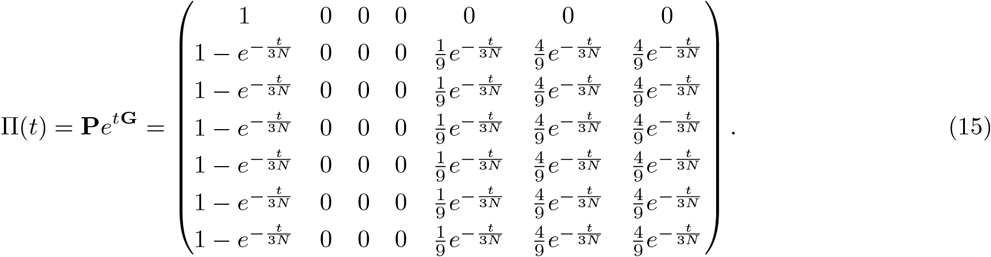

Here, examining the first column of the matrix in Eq. 15—representing transitions to coalescence—we can see that two lineages within an individual (state 1), within a mating pair (state 2_0_), or in in two separate mating pairs (states 3, 4, and 5) have equal coalescence times. In fact, as coalescence times are unaffected by patrilateral-parallel first-cousin consanguinity, they accord with the coalescence time distribution for a population of size 3*N* haploid individuals. Using the same random variables from the sibling case (where *U* now represents 2_0_), we can extract the cumulative distribution functions of coalescence times from the first column of the matrix Π(*t*):

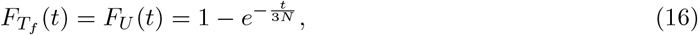

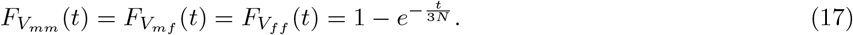

For each of the five random random variables, the time to coalescence for two lineages is distributed as an exponential random variable with rate 1*/*(3*N*). The mean of these distributions—the reciprocal of the coalescence rate—is 3*N*, matching the limiting means obtained by first-step analysis in Eqs. 28–32 of Cotter *et al*. (2021).

#### 3.2.2 Patrilateral cross

For the patrilateral-cross case, a union occurs between a male and his father’s sister’s daughter. As with the parallel case, there is no way for the X-chromosomal lineages in the first-cousin mating pair to have originated from a shared ancestor. We obtain the exact same transition probabilities from state 2_1_ and the same transition matrix (Eq. 12). The coalescence times for the patrilateral-cross case are the same as in the patrilateral-parallel case.

#### 3.2.3 Matrilateral parallel

In the matrilateral parallel case, a union occurs between a male and his mother’s sister’s daughter. With probability *c*_1_*/*2, two lineages in state 2_1_ trace back to the shared grandparental pair. The lineages in state 2_1_ coalesce with probability 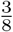 (state 0), they are in the shared grandmother with probability 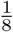 (state 1), and they are in opposite individuals of the grandparental mating pair with probability 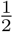 (state 2_0_).

With probability *c*_1_*/*2, two lineages in state 2_1_ do not trace back to the shared grandparental pair. Conditional on not tracing to this pair, they are in a male and a female (state 4) or two females (state 5), each with probability 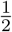. Finally, with probability 1 − *c*_1_, the two lineages are not ancestral to a consanguineous mating pair; they then follow the same pattern as in the patrilateral-parallel case. Combining the cases gives the transition matrix,

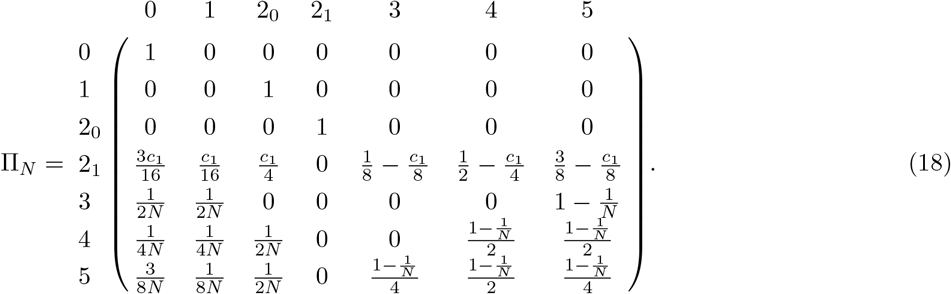

As before, we decompose this matrix into “fast” and “slow” transitions (Eq. 1):

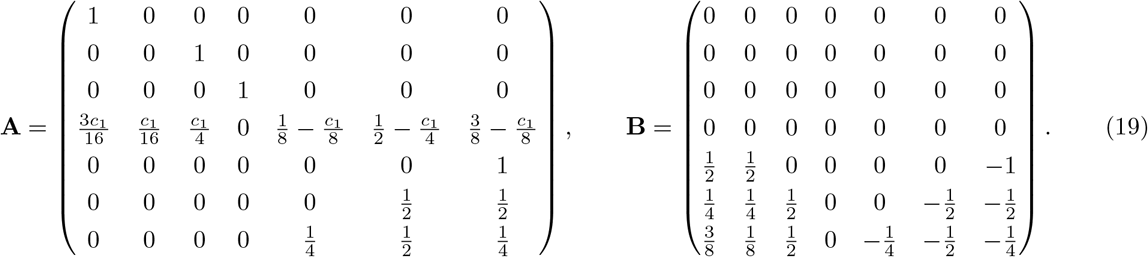

We next solve for the limiting distribution of the fast matrix **A** using the method of Appendix A:

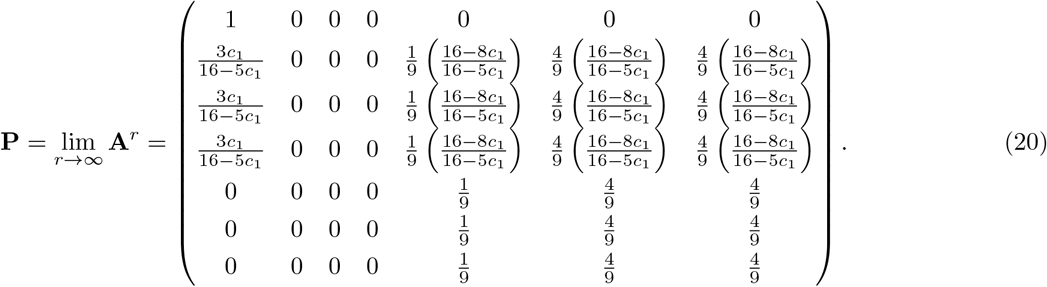

Finally, recalling **G** = **PBP**, we solve for the matrix exponential *e*^*t***G**^ using the method of Appendix B. We then solve for the continuous-time process Π(*t*) via Eq. 3, converting *t* back to units of *N* generations:

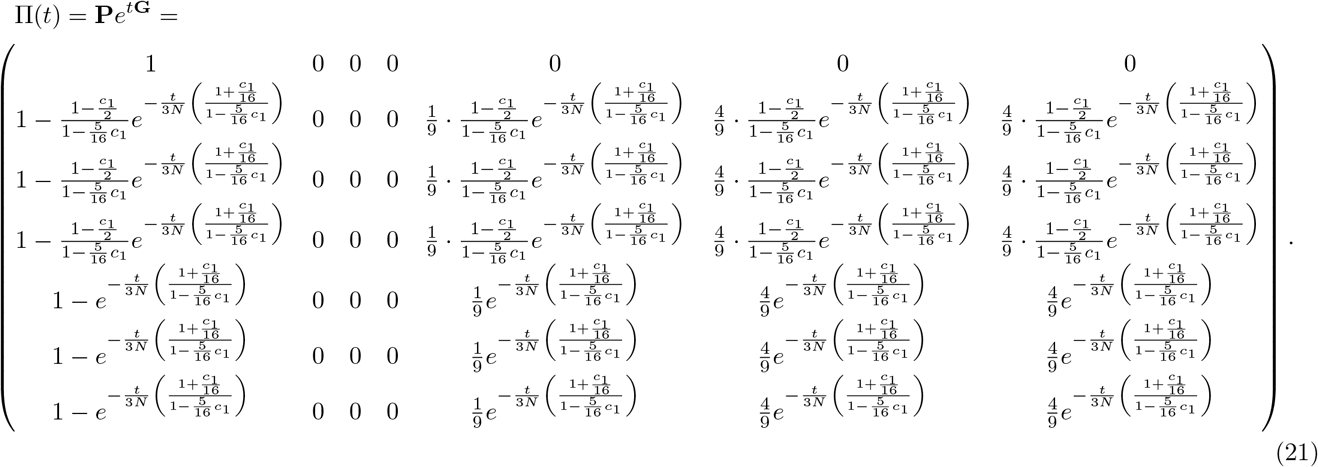

We are concerned with transitions from each of the various states to coalescence (state 0). The first column of Π(*t*) gives the limiting cumulative distribution functions for the time to the most recent common ancestor for two lineages *within* an individual (state 1) and two lineages *between* individuals (states 3, 4 and 5):

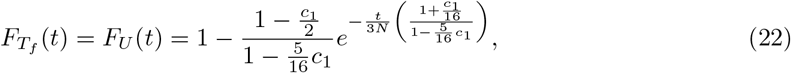

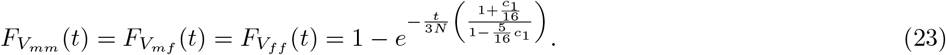

To compute expectations, recalling that for 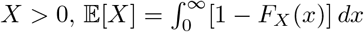, we find

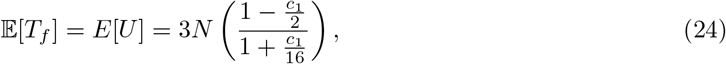

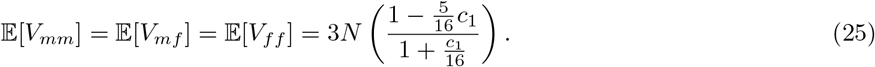

Eqs. 24 and 25 are the same as Eqs. 39 and 40 from Cotter *et al*. (2021). Eqs. 22 and 23 are plotted in Figure 5.

**Figure 5:**
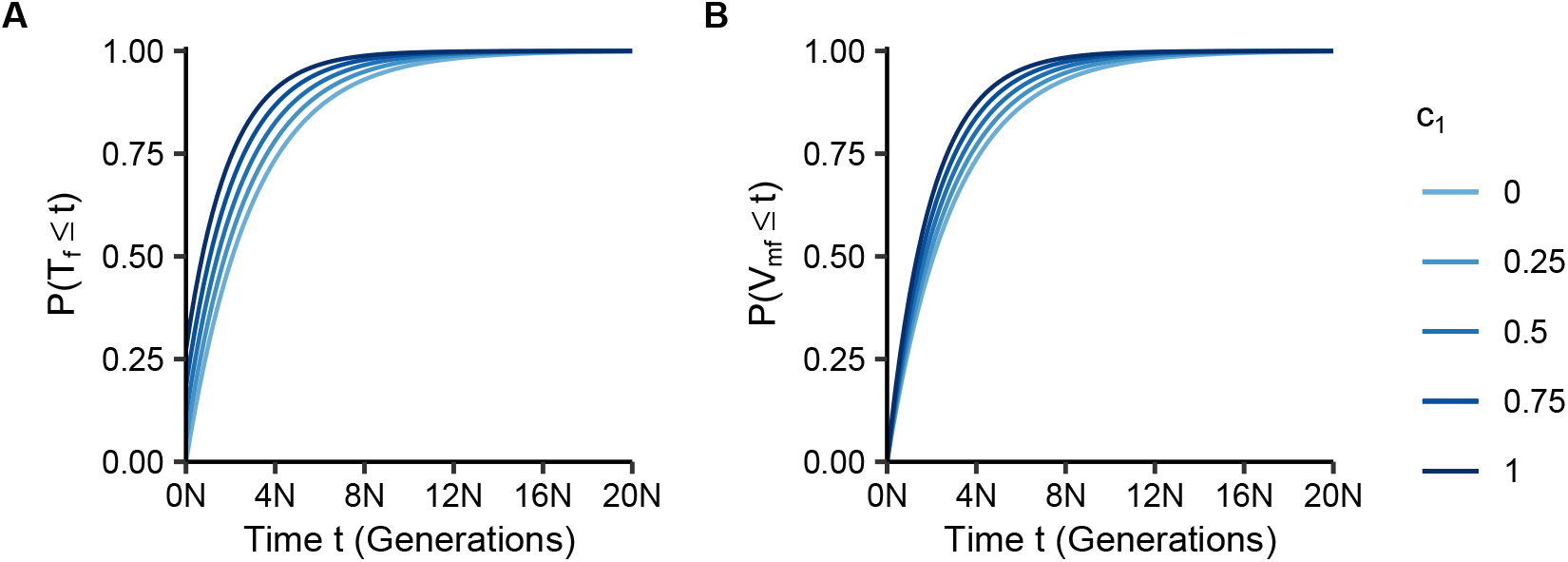
Cumulative distributions of coalescence times within (*T*_*f*_) and between (*V*_*mf*_) individuals as functions of the number of generations *t* and the fraction of matrilateral-parallel mating pairs *c*_1_. **(A)** Within individuals, *P* (*T*_*f*_ ≤ *t*), Eq. 22. **(B)** Between individuals, *P* (*V*_*mf*_ ≤ *t*), Eq. 23.

#### 3.2.4 Matrilateral cross

In the matrilateral-cross case, a union occurs between a male and his mother’s brother’s daughter. This case is similar to the matrilateral-parallel case. With probability *c*_1_*/*2, two lineages in state 2_1_ trace to the shared grandparental pair. They coalesce with probability 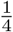 (state 0), they are in the shared grandmother with probability 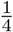 (state 1), and they are in opposite individuals of the grandparental mating pair with probability 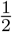 (state 2_0_).

With probability *c*_1_*/*2, two lineages in state 2_1_ do not trace to the shared grandparental pair. Conditional on the lineages not both tracing to the shared grandparental pair, they are in two males (state 3), a male and a female (state 4) or two females (state 5), with probabilities 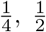, and 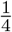, respectively. Finally, with probability 1 − *c*_1_, two lineages are not ancestral to a consanguineous mating pair. In this case, they follow the same pattern as enumerated for the patrilateral-parallel case. The transition matrix is

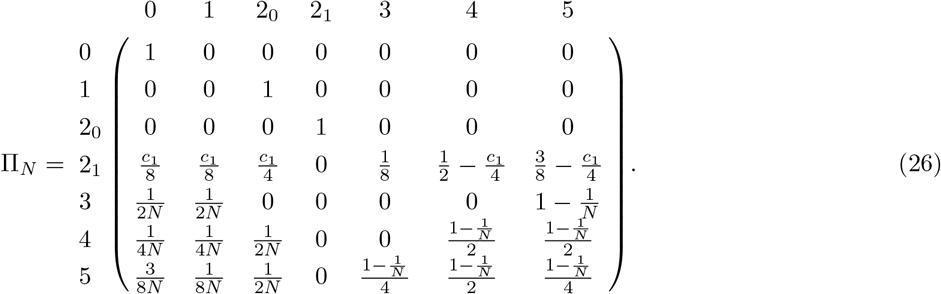

We separate the “fast” and “slow” transitions as before (Eq. 1):

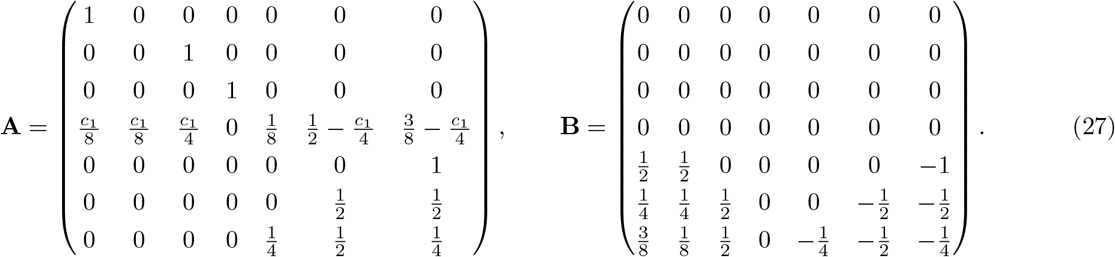

Using the method of Appendix A, we solve for the stationary distribution of the “fast” process:

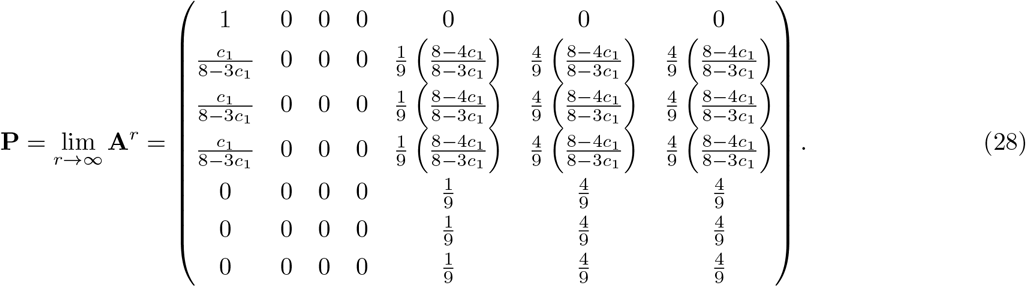

As before, using **G** = **PBP**, we calculate the matrix exponential, *e*^*t***G**^, using the method of Appendix B. We then obtain Π(*t*) from Eq. 3, converting *t* back to units of *N* generations:

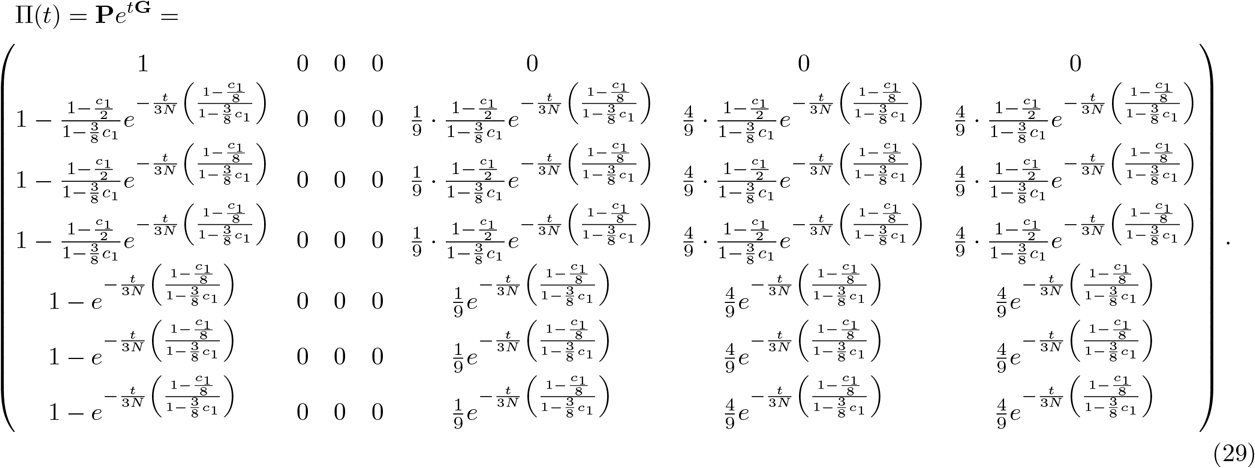

We extract the cumulative distribution functions from the first column of the matrix, finding

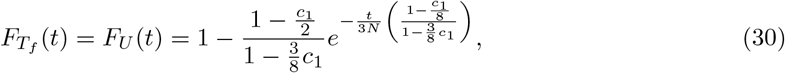

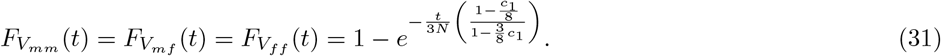

Solving for the expectations of these distributions, recalling that for 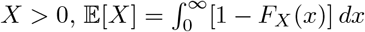, we find

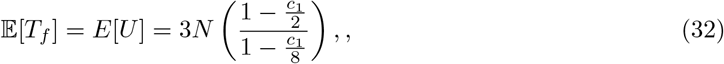

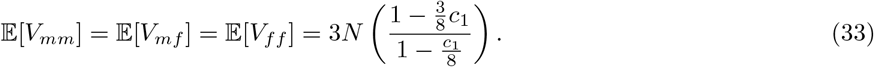

Eqs. 32 and 33 are the same as Eqs. 47 and 48 from Cotter *et al*. (2021). Eqs. 30 and 31 are plotted in Figure 6.

**Figure 6:**
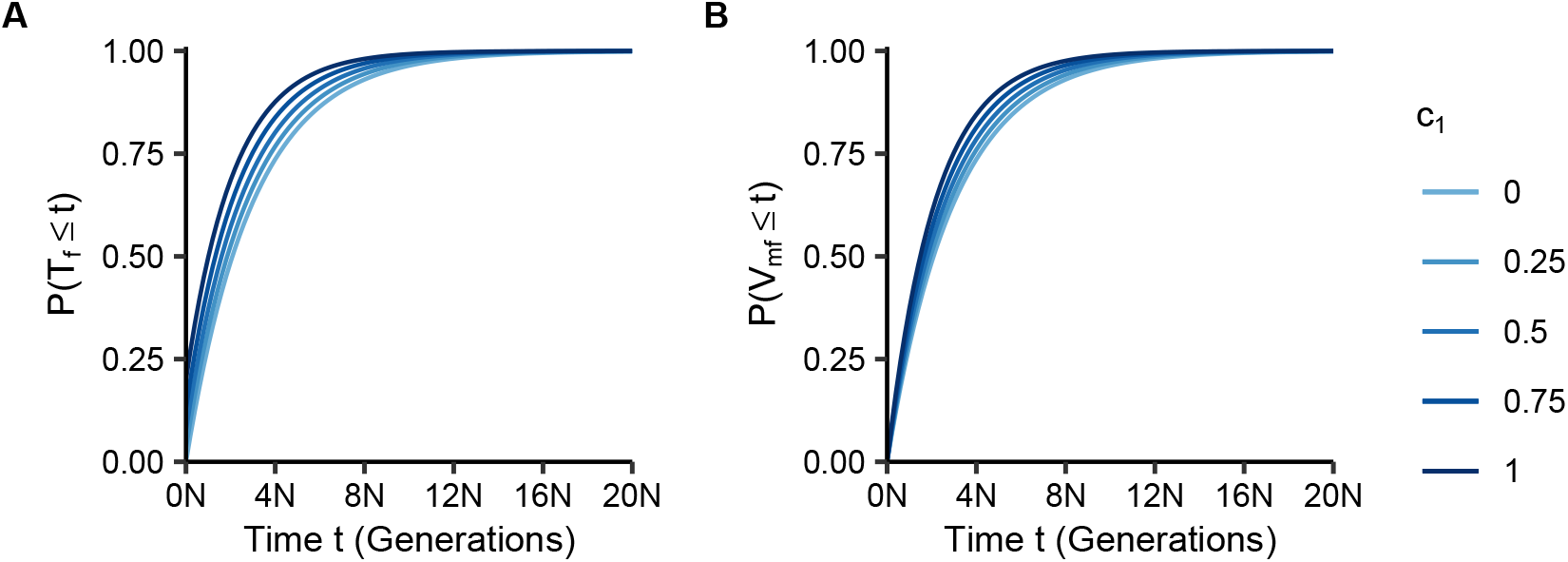
Cumulative distributions of coalescence times within (*T*_*f*_) and between (*V*_*mf*_) individuals as functions of the number of generations *t* and the fraction of matrilateral-cross mating pairs *c*_1_. **(A)** Within individuals, *P* (*T*_*f*_ ≤ *t*), Eq. 30. **(B)** Between individuals, *P* (*V*_*mf*_ ≤ *t*), Eq. 31.

#### 3.2.5 Bilateral parallel

Having considered the four possible types of first-cousin consanguinity, we can also consider the two bilateral cases, in which a mating pair are cousins through both sets of grandparents. In bilateral-parallel first-cousin consanguinity, a union occurs between a male and a female who is both his mother’s sister’s daughter *and* his father’s brother’s daughter. We can consider this case to be a combination of the matrilateral-parallel case and the patrilateral-parallel case. In state 2_1_, when the two lineages are ancestral to a bilateral-parallel mating pair, the male’s lineage must transition through his mother because he cannot inherit an X chromosome from his father. Because there is no way for the lineages to transition through the patrilateral-parallel grandparental pair, the transitions in state 2_1_ follow from the transitions for a matrilateral-parallel pair only. In the case of bilateral-parallel first-cousin consanguinity, the transition matrix thus has the form given for matrilateral-parallel first-cousin consanguinity in Eq. 18. The bilateral-parallel case thus also shares the same cumulative distribution functions given in Eqs. 22 and 23.

#### 3.2.6 Bilateral cross

Bilateral-cross first-cousin consanguinity occurs when a male shares a union with a female who is both his father’s sister’s daughter and his mother’s brother’s daughter. This case can be considered to be a combination of matrilateral-cross and patrilateral-cross first-cousin consanguinity. The ancestral lineages cannot travel through the patrilateral-cross pair, and the transitions follow those for matrilateral-cross consanguinity. The transition matrix (Eq. 26) and cumulative distribution functions (Eqs. 30 and 31) follow similarly.

#### 3.2.7 Mixture of first-cousin mating types

We next examine a population that possesses a mixture of all four unilateral first-cousin mating types. To determine the transition matrix, it suffices to determine the transition probabilities from state 2_1_.

Recall that two lineages in state 2_1_ are in two individuals ancestral to a mating pair that might or might not be consanguineous. With probability *c*_*pp*_, this mating pair is a patrilateral-parallel first-cousin pair, with probability *c*_*pc*_ it is a patrilateral-cross first-cousin pair, with probability *c*_*mp*_ it is a matrilateral-parallel first-cousin pair, and with probability *c*_*mc*_ it is a matrilateral-cross first-cousin pair. If the mating pair is a first-cousin pair of a particular one of the four types, then transitions out of state 2_1_ will match those derived for the associated case.

We can view the transition probabilities out of state 2_1_ as a weighted combination of the transitions that each of these first-cousin cases makes when considered on its own. For example, in the case of coalescence (transition to state 0), two lineages in state 2_1_ coalesce with probability 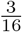 for a matrilateral-parallel first-cousin pair (rate *c*_*mp*_) and 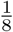 for a matrilateral-cross first-cousin pair (rate *c*_*mc*_). Because patrilateral-parallel and patrilateral-cross consanguinity do not affect transitions from state 2_1_, corresponding rates *c*_*pp*_ and *c*_*pc*_ do not influence the transition probability to state 0. Combining all four cases, the transition probability from state 2_1_ to state 0 is 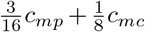. For transitions from state 2_1_ to states 0, 1, and 2_0_, the probabilities are obtained by summing corresponding terms in the matrices for the various types of unilateral first-cousin mating (Eqs. 12, 18, and 26).

For the transitions from state 2_1_ to states 3, 4, and 5 (two lineages between individuals), consanguinity acts to reduce the probabilities. The probabilities in the case of patrilateral parallel consanguinity (Eq. 12) represent a null effect of no consanguinity. The *c*_*mp*_ and *c*_*mc*_ terms (Eqs. 18 and 26) reduce the probabilities of transitioning to states 3, 4, and 5 (while inflating the 0, 1, and 2_0_ transitions). For state 3, for example, the null transition probability is 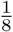. Matrilateral-parallel consanguinity reduces this transition probability by *c*_*mp*_*/*8, giving a combined transition probability of 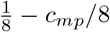 matrilateral-cross consanguinity has no effect on this transition.

We proceed similarly to combine the remaining transition probabilities from the four unilateral first-cousin mating types to produce the transitions for state 2_1_. The transition matrix is

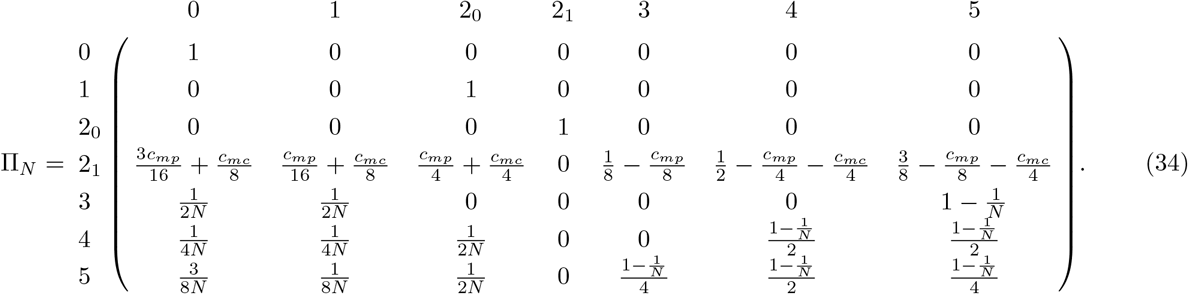

Matrices **A** and **B** follow from Eq. 1 and take the same form as those given for the matrilateral cases with state 2_1_ in matrix **A** (Eqs. 19 and 27), now adopting the new combinations of transition probabilities. We solve for the stationary distribution of the “fast” transitions using the method of Appendix A:

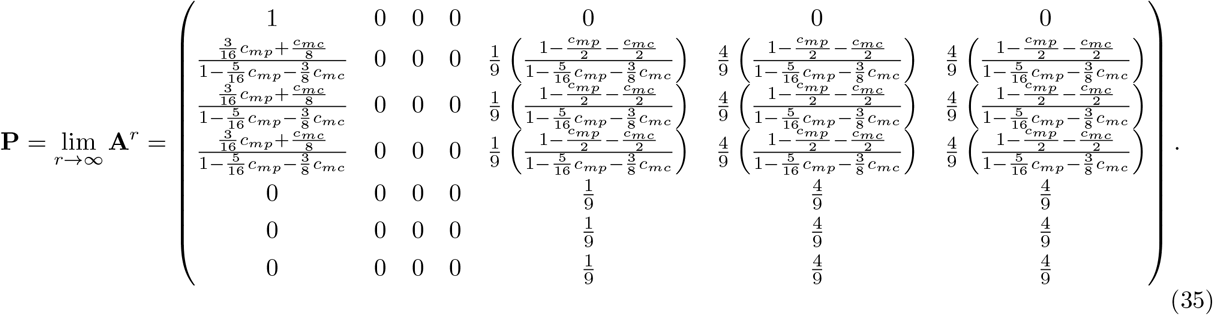

Once again, using **G** = **PBP**, we obtain the matrix exponential, *e*^*t***G**^, using the method of Appendix B. We then compute Π(*t*) with Eq. 3, converting *t* back into units of *N* generations. The resulting matrix is structured in such a way that we can write:

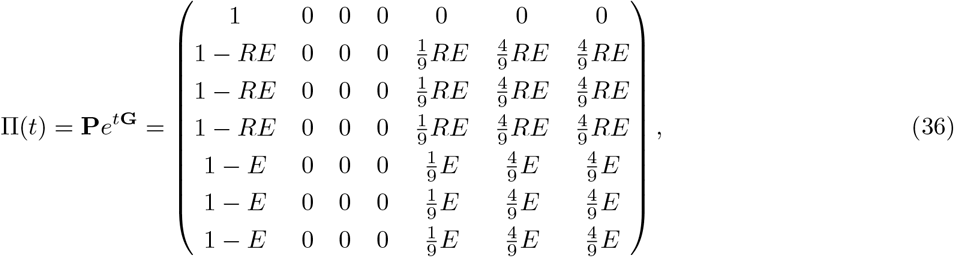

Where

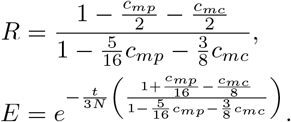

In the matrix in Eq. 36, the first column represents transitions to coalescence. We extract from this column the cumulative distribution functions for time to coalescence for two lineages *within* an individual (state 1) and two lineages *between* individuals (states 3, 4, and 5):

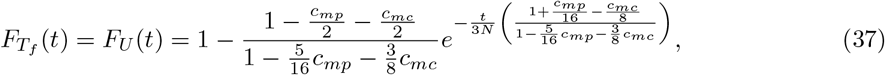

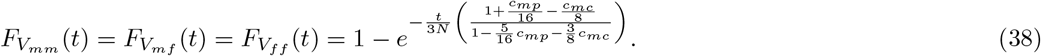

Solving for the expectations of these distributions, recalling that for 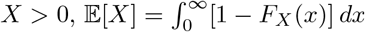, we find

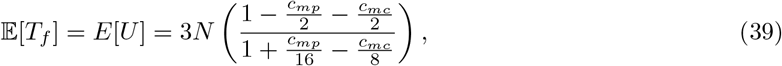

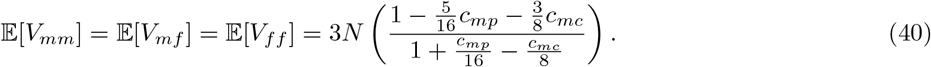

### 3.3 Comparisons

#### 3.3.1 Limiting distribution versus exact distribution

Under the mixture model, to see how well the limiting distribution of coalescence times approximates the exact distribution, we perform simulations. In particular, for fixed values of the number of mating pairs *N* and rates of matrilateral-parallel (*c*_*mp*_) and matrilateral-cross (*c*_*mc*_) first-cousin mating, we simulate 10,000 realizations of the Markov chain in Eq. 34 to produce an empirical cumulative distribution function (CDF) of coalescence times for lineage pairs *within* and *between* individuals. This procedure amounts to simulating a distribution of the time to the most recent common ancestor (the time it takes to hit state 0) starting in either state 1 (within an individual) or state 4 (between individuals).

Figure 7 plots the simulated empirical CDFs alongside the limiting CDFs presented in Eqs. 37 and 38. Conducting these simulations for different values of the number of mating pairs *N*, we see that as *N* increases, the limiting distribution functions (Eqs. 37 and 38) closely approximate the simulated, empirical distributions.

**Figure 7:**
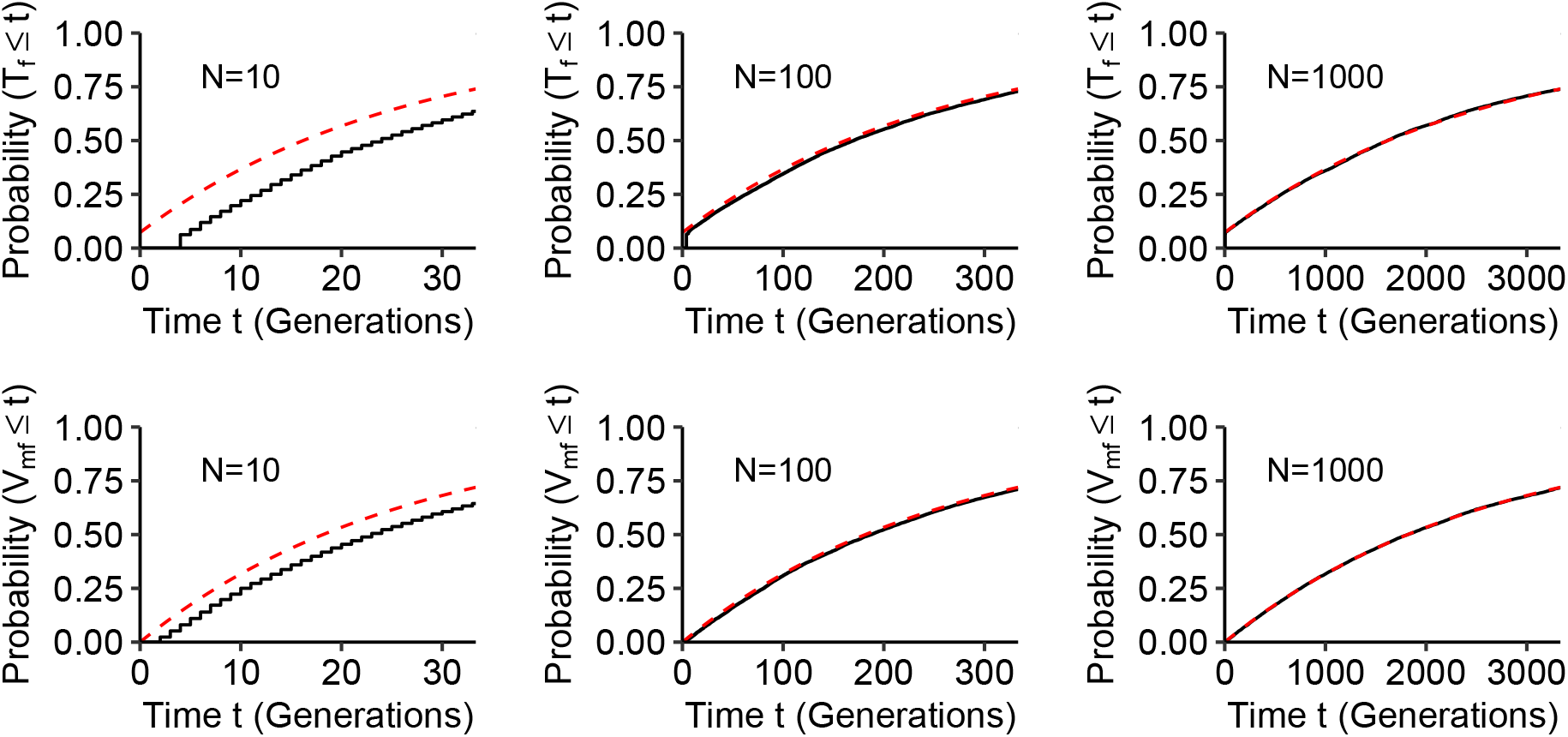
Cumulative distribution functions (CDFs) of coalescence times in a model with a mixture of types of consanguinity. The Markov chain is given in Eq. 34; we consider the case of *c*_*mp*_ = 0.2 and *c*_*mc*_ = 0.2 with each of three values for the number of mating pairs *N*. Dashed lines represent the limiting CDFs in Eqs. 37 and 38, and solid lines represent the simulated CDFs from 10,000 observations of the first-cousin mixture model (as described by the Markov chain in Eq. 34).

#### 3.3.2 X chromosome versus autosomes

Each of the limiting distributions for coalescence times for lineages from separate mating pairs, both for single types of first-cousin consanguinity and for a superposition of multiple types, possesses a particular structure: an exponential CDF whose rate is the product of the population size and a reduction by a factor that accounts for consanguinity. We now examine these limiting CDFs for the X chromosome in relation to corresponding CDFs for autosomes. The autosomal coalescence time distributions under first-cousin consanguinity are obtained in Appendix C as a special case of the *n*th cousin mating model of Severson *et al*. (2019). Here, we calculate the ratio of the expected time to coalescence for the X chromosome (Eqs. 39 and 40) and for autosomes (Eqs. C4 and C5) within and between individuals, respectively, as we vary rates of matrilateral and patrilateral consanguinity (Figure 8).

**Figure 8:**
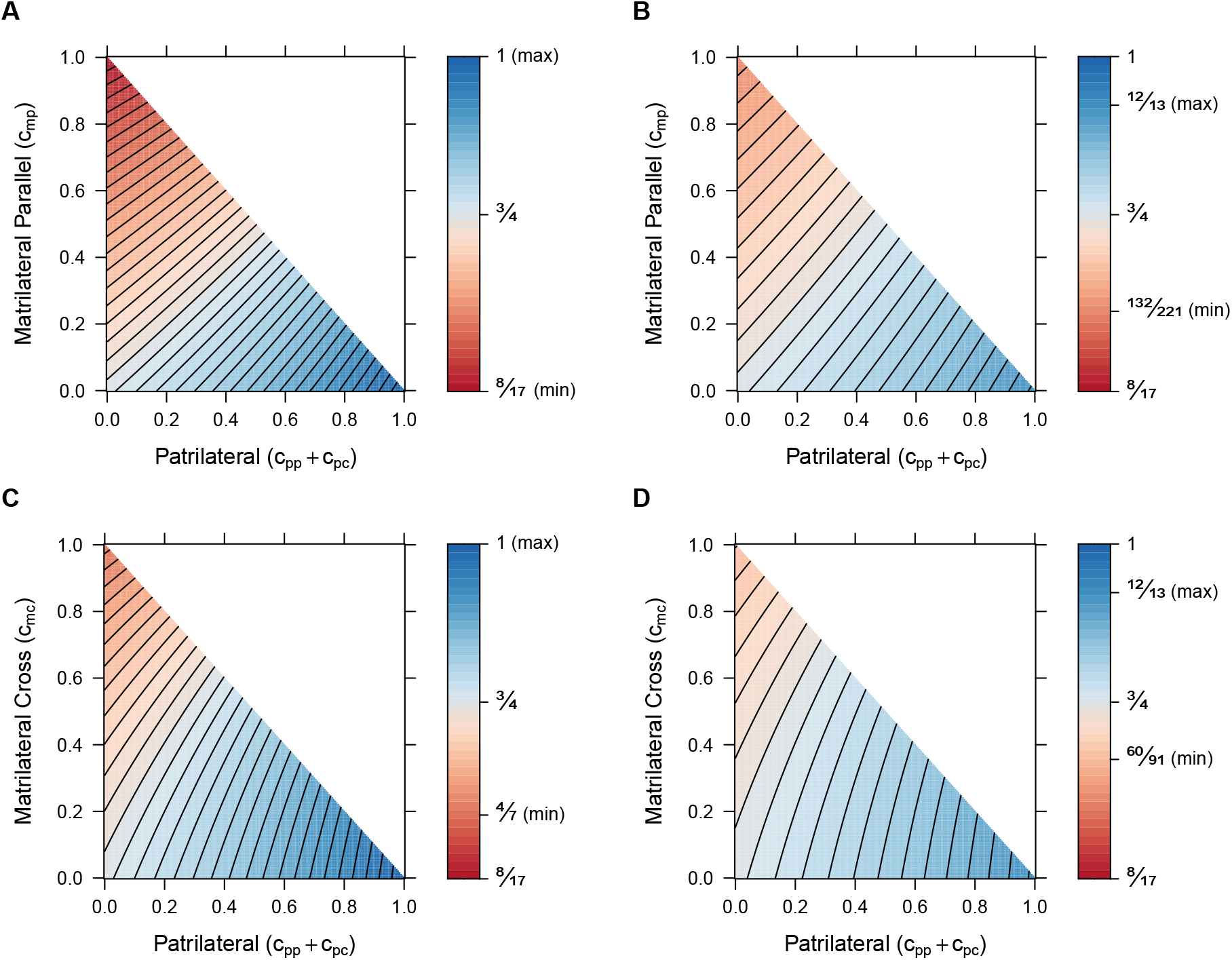
Ratios of X-chromosomal and autosomal mean coalescence times. Each point represents a ratio of coalescence times for a specified mixture of two types of consanguinity, depicted on the x and y axes. (A) Within individuals, matrilateral parallel and patrilateral consanguinity (Eq. 39/Eq. C4). (B) Between individuals, matrilateral parallel and patrilateral consanguinity (Eq. 40/Eq. C5). (C) Within individuals, matrilateral cross and patrilateral consanguinity (Eq. 39/Eq. C4). (D) Between individuals, matrilateral cross and patrilateral consanguinity (Eq. 40/Eq. C5). In each panel, the minimal ratio is indicated (obtained by setting matrilateral consanguinity to 1 and patrilateral consanguinity to 0), as is the maximum (obtained by setting matrilateral consanguinity to 0 and patrilateral consanguinity to 1). The value 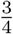 occurs with no consanguinity, located at the origin in each panel. Values *greater* than 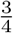 appear in blue, indicating combinations of parameter values that bring expected X chromosomal coalescence times closer to expected autosomal coalescence times. Values that reduce X chromosomal coalescence times to a greater extent than on autosomes, thereby shifting the ratio less than 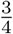, appear in red.

We first consider the ratio of expected coalescence times on the X chromosome relative to the autosomes for pairs of lineages within individuals (Eq. 39/Eq. C4) as a function of patrilateral (*c*_*pp*_ + *c*_*pc*_) and matrilateral-parallel (*c*_*mp*_) consanguinity (Figure 8A). Because the expected coalescence time for two lineages on the X chromosome is a function of 3*N* and the corresponding autosomal mean depends on 4*N*, in the absence of consanguinity, the null value of the ratio is 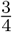. The ratio achieves its minimum value of 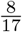, with a stronger effect of consanguinity in reducing X-chromosomal coalescence times relative to autosomal coalescence times, when we set *c*_*mp*_ to 1. It achieves its maximum value of 1, increasing X-chromosomal coalescence times compared to autosomal coalescence times, when instead we set *c*_*pp*_ + *c*_*pc*_ to 1 (Figure 8A).

For the X:A ratio of between-individual expected coalescence times (Eq. 40/Eq. C5) as a function of patrilateral (*c*_*pp*_ + *c*_*pc*_) and matrilateral-parallel (*c*_*mp*_) consanguinity (Figure 8B), the minimum and maximum values differ less than for the within-individual case. The minimum exceeds 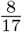, equaling 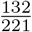, and is again reached at *c*_*mp*_ = 1. The maximum is less than 1, equaling 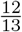, and is reached at *c*_*pp*_ + *c*_*pc*_ = 1. The minimum and maximum are less extreme than in the within-individual case, as consanguinity has less of an effect on reducing the expected coalescence times in the between-individual case, both for the X chromosome and for the autosomes.

We next examine the X:A coalescence time ratio within individuals (Eq. 39/Eq. C4) as a function of patrilateral (*c*_*pp*_ + *c*_*pc*_) and matrilateral-cross (*c*_*mc*_) consanguinity (Figure 8C). The minimal ratio is slightly larger than in the matrilateral-parallel case, equaling 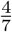 at *c*_*mc*_ = 1. The maximum occurs at 1, the same value as the corresponding case with matrilateral-parallel in place of matrilateral-cross consanguinity, when *c*_*pp*_ + *c*_*pc*_ = 1. The slightly reduced range of values (i.e., the greater minimum) traces to the fact that the effect of matrilateral-cross consanguinity on X-chromosomal coalescence times is slightly weaker, producing a weaker reduction in coalescence times, than that of matrilateral-parallel consanguinity.

Finally, we analyze the X:A coalescence time ratio between individuals (Eq. 40/Eq. C5) as a function of patrilateral (*c*_*pp*_ + *c*_*pc*_) and matrilateral-cross (*c*_*mc*_) consanguinity (Figure 8D). The minimum occurs at *c*_*mc*_ = 1, equaling 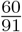. As in the corresponding matrilateral-parallel case, the maximum, achieved at *c*_*pp*_ + *c*_*pc*_ = 1, is 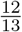. As was seen within individuals, the range of permissible values is reduced relative to the matrilateral-parallel case, owing again to the weaker effect of matrilateral-cross consanguinity on X-chromosomal coalescence times.

## 4 Discussion

Extending our previous work on mean coalescence times on the X-chromosome in a consanguinity model, we have derived large-*N* limiting distributions for within-individual and between-individual X-chromosomal coalescence times under various types of first-cousin consanguinity. For between-individual coalescence times, each limiting distribution is exponential with a rate equal to the product of the number of X chromosomes and a reduction factor due to consanguinity (Eqs. 17, 23, and 31). Limiting distributions of within-individual coalescence times each have a point mass corresponding to instantaneous coalescence, and conditional on not coalescing instantaneously, are exponential (Eqs. 16, 22, and 30). These patterns also hold for limiting distributions of pairwise coalescence times for a model with a mixture of types of first-cousin consanguinity (Eqs. 37 and 38); in simulations, the limiting distributions under this superposition agree with exact distributions from the Markov chain (Eq. 34, Figure 7).

Our limiting distribution results can inform comparisons of the X chromosome with autosomes. The four types of first-cousin consanguinity have identical effects on the autosomes but vary in their effect on the X chromosome. Hence, a comparison of coalescence time distributions for the X chromosome and autosomes can be informative about features of consanguinity. Our results (Eqs. 37 and 38) directly show the effect of different rates and types of consanguinity on the distribution of X-chromosomal coalescence times. For example, increasing matrilateral-parallel and matrilateral-cross consanguinity decreases the ratio of X and autosomal mean coalescence times; increasing patrilateral-parallel and patrilateral-cross first-cousin consanguinity increases this ratio (Figure 8).

Consanguinity and other preferences for mate choice vary across populations, often depending on cultural norms for certain types of consanguinity over others (Bittles, 2012). Because we have found that the different types of first-cousin consanguinity generate an observable effect on X chromosomal coalescence times, it is possible that features of coalescence times can be compared across populations to assess signatures of the different types of consanguinity. Such assessments can potentially capitalize on the inverse relationship between coalescence times and genomic sharing (Palamara *et al*., 2012; Carmi *et al*., 2014; Browning and Browning, 2015) to use genomic sharing patterns to uncover features of consanguinity (Arciero *et al*., 2021).

One limitation of our approach is that in formulating our model, we have disregarded higher-order consanguinity. While we have explicitly modeled first-cousin mating pairs, we have ignored the possibility that a pair has more distant consanguinity that is not captured in the model. It may be possible, however, to allow for such possibilities by incorporating into the *n*th cousin framework of Severson *et al*. (2021) sex-specific varieties of consanguinity at different levels of relationship.

## Acknowledgments

We acknowledge support from United States–Israel Binational Science Foundation grant 2017024, NIH grant R01 HG005855, and NSF Graduate Research Fellowships to DJC and ALS.

## Appendix A: Stationary distribution of the fast transition matrix

In this appendix, we solve for the stationary distribution of the “fast” transition matrix **A** in the case of sib mating on the X chromosome. The same approach is also applied in the main text to obtain the stationary distribution of the fast transition matrix in other models.

First, we permute the states to rewrite matrix **A** in a canonical form. The matrix **A** in Eq. 5 has one absorbing state (state 0) and a closed communication class *C*_1_ = *{*3, 4, 5*}*. For simplicity, we write the sib mating probability *c*_0_ as *c*. We rearrange the matrix to take the form

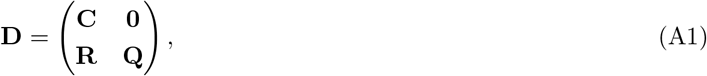

listing the recurrent states before the transient states. Thus, square matrix **C** includes transitions between recurrent states (i.e., absorbing states and closed communication classes), and square matrix **Q** includes transitions between transient states. Matrix **R** includes transitions from the transient states to the recurrent states. For matrix **A** in Eq. 5, the recurrent states are state 0 (absorbing) and states 3, 4, and 5 (closed communication class *C*_1_). The transient states are states 1 and 2. Permuting the matrix **A** to order the states 0, 3, 4, 5, 1, 2, we write

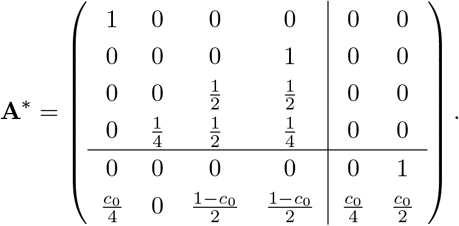

We treat the closed communication class *C*_1_ as a single absorbing state because any transitions made into *C*_1_ transition infinitely often among the states it contains. We rewrite the transition matrix for the resulting Markov chain by collapsing the columns and rows corresponding to the states in *C*_1_. **A**^*∗*^ becomes

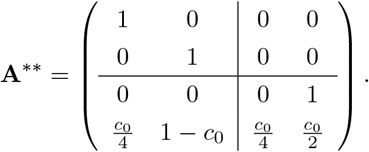

Matrix **A**^*∗∗*^ now has the form in Eq. A1, with 2 *×* 2 submatrices and **C** as the identity matrix. Given a matrix in canonical form (Eq. A1 where **C** is the identity), the stationary distribution is given by

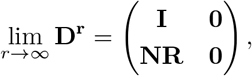

where *N* is the fundamental matrix **N** = (**I** − **Q**)^−1^ and **I** is the identity matrix (Kemeny and Snell, 1983, 3.3.7). The matrix **NR** defines for each pair consisting of a transient state and a recurrent state, the probability that from the transient state, the process reaches the recurrent state. For matrix **A**^*∗∗*^, we have

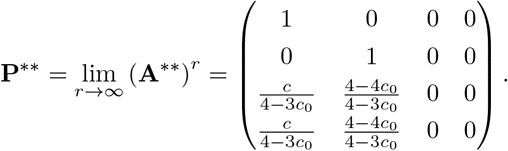

To recover the stationary distribution of **A**^*∗*^, we expand the absorbing state for the closed communication class *C*_1_, replacing it with the stationary distribution for the irreducible 3 *×* 3 matrix associated with the class. We then weight the transient transition probabilities in **NR** by this stationary distribution.

In other words, **NR** now gives, for each pair consisting of a transient and a recurrent state, the probability of the associated transition. Expanding the absorbing state for the closed communication class *C*_1_, we get

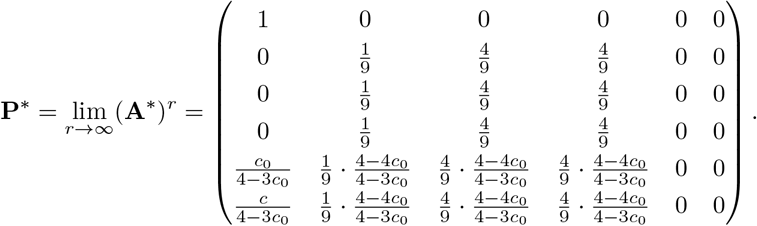

Finally, we permute **P**^*∗*^ to recover **P** (Eq. 6).

## Appendix B: The matrix exponential *e*^*t*G^

In this appendix, we obtain the matrix exponential, *e*^*t***G**^, which is needed in calculating the large-*N* limit, Π(*t*) = **P***e*^*t***G**^. The computations in this appendix are specific to sib mating on the X chromosome, but the same method can be applied to obtain the matrix exponential in the other models.

We first obtain the generator matrix from Eqs. 5 and 6:

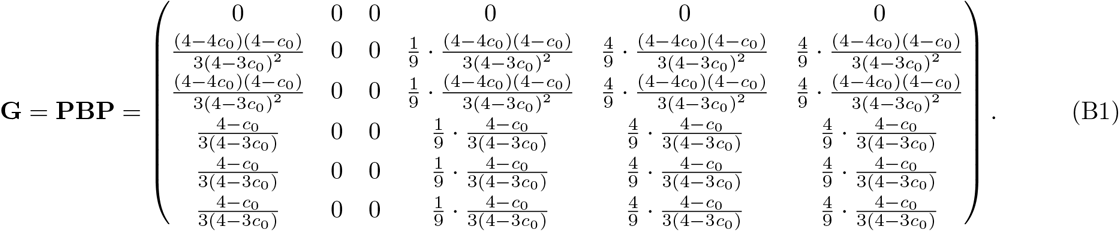

The generator matrix, **G**, has nonzero entries in the columns for state 0 and states 3, 4, and 5. It has the property

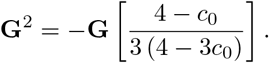

For the constant *k* = −(4 − *c*_0_)*/*[3 (4 − 3*c*_0_)], we can then recursively write

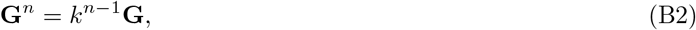

The matrix exponential, 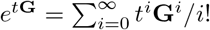, then equals

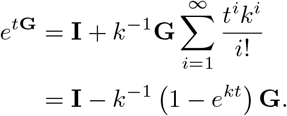

Converting *t* into units of *N* generations and multiplying by **P** (Eq. 6), we obtain **P***e*^*t***G**^ as in Eq. 7.

For each model studied, for the associated generator matrix **G**, the corresponding quantity *k* that satisfies Eq. B2 appears in Table B1.

## Appendix C: Limiting distribution of autosomal coalescence times for first-cousin mating

Equation 46 of Severson *et al*. (2021) gives a limiting distribution of autosomal coalescence times for a model with a superposition of levels of cousin mating, up to *n*th cousins. In order to recover first-cousin mating on the autosomes to compare to our X-chromosomal results, we use the special case of this *n*th cousin model, where the rate of sibling mating *c*_0_ is 0 and the rate of first-cousin mating is *c*_1_, stopping at first cousins. This special case produces the following transition matrix where state 0 is still coalescence, state 1 is two lineages in an individual, state 2_0_ is two lineages in opposite individuals of a mating pair, state 2_1_ is two lineages in two individuals one generation ancestral to a mating pair, and state 3 is two lineages in two individuals in different mating pairs:

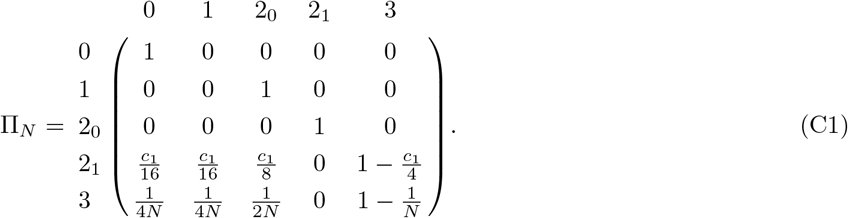

Note here that there is no need to use a two-sex model, as for autosomes, states referring to two males, a male and a female, and two females simply collapse into the combined state 3. No new information is gained for the autosomes when separating these states. Using Eq. 1, we split the transition matrix into fast and slow processes:

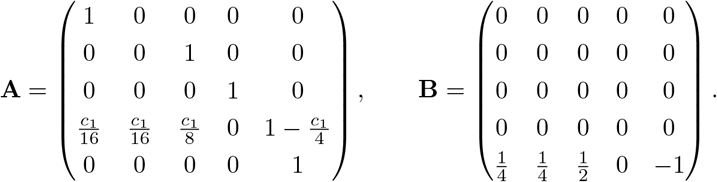

We solve for the stationary distribution of the fast matrix using the method in Appendix A (simpler here by a single absorbing state for two lineages between individuals rather than a closed communication class):

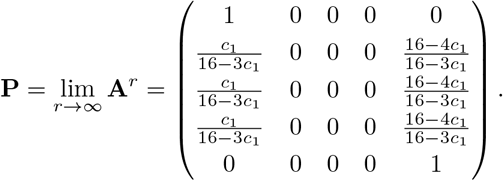

Using **G** = **PBP**, we obtain the matrix exponential *e*^*t***G**^ using the method of Appendix B. We then compute Π(*t*) via Eq. 3, converting *t* back into units of *N* generations:

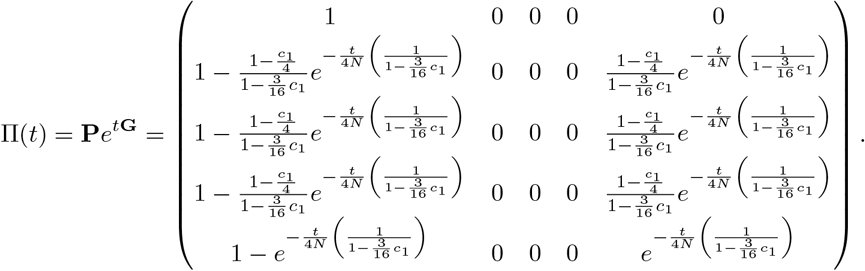

We extract from the first column of this matrix the cumulative distribution functions for two lineages starting in state 1 (within an individual) and state 3 (between individuals):

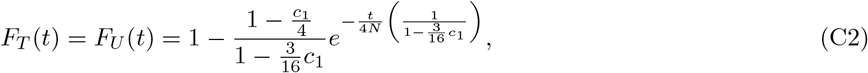

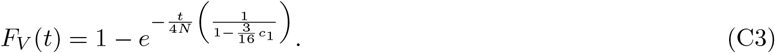

Severson *et al*. (2021) showed that the limiting distribution for *n*th cousin mating is given by their Eqs. 47 and 48:

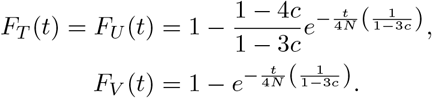

In the special case where we only have first-cousin mating, we replace their *c* term with *c*_1_*/*16 and recover Eqs. C2 and C3, respectively.

For the expectations of these distributions, by 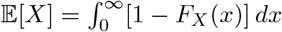, for *X* > 0, we find

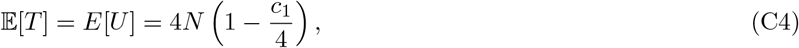

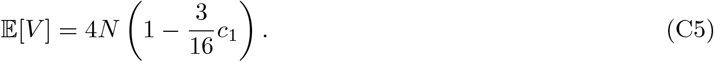

Eqs. C4 and C5, obtained from the limiting distribution, accord with the large-*N* limit of Eqs. 8 and 10 from Severson *et al*. (2019), in which they were calculated via first-step analysis.

---

**Table B1:** Constants used in matrix exponentiation for consanguinity models.

| Type of consanguineous mating | Chromosome | Section | Quantity $k$ satisfying $G^n = k^{n-1}G$ for generator matrix $G$ (Eq. B2) |
| --- | --- | --- | --- |
| Sibling | X | 3.1 | $-\frac{4-c_0}{3(4-3c_0)}$ |
| Patrilateral-parallel first-cousin | X | 3.2.1 | $-\frac{1}{3}$ |
| Patrilateral-cross first-cousin | X | 3.2.2 | $-\frac{1}{3}$ |
| Matrilateral-parallel first-cousin | X | 3.2.3 | $-\frac{16+c_1}{3(16-5c_1)}$ |
| Matrilateral-cross first-cousin | X | 3.2.4 | $-\frac{8-c_1}{3(8-3c_1)}$ |
| Bilateral-parallel first-cousin | X | 3.2.5 | $-\frac{16+c_1}{3(16-5c_1)}$ |
| Bilateral-cross first-cousin | X | 3.2.6 | $-\frac{8-c_1}{3(8-3c_1)}$ |
| Superposition of unilateral first-cousin | X | 3.2.7 | $-\frac{16+c_{mp}-2c_{mc}}{3(16-5c_{mp}-6c_{mc})}$ |
| First-cousin | Autosomes | Appendix C | $-\frac{4}{16-3c_1}$ |

Note that $c_{mp}$ and $c_{mc}$ in Section 3.2.7 have the same meaning as $c_1$ in Sections 3.2.3 and 3.2.4, respectively.

